# Bravais Lattice Sampling: Geometry-Guided Sparse Probing for Connected-Component Detection in 3D Discretized Spaces

**DOI:** 10.64898/2026.09.02.748950

**Authors:** Francisco Carrascoza

## Abstract

We introduce Bravais Lattice Sampling (BLS), a two-phase method for detecting connected high-density regions in three-dimensional space. BLS places probe sites on a Bravais lattice scaled to the expected nearest-neighbour distance *d*_NN_ of the target structures, then recovers cluster boundaries by depth-first expansion seeded only from occupied probes, replacing the exhaustive raster scan that conventional connected-component labelling uses to discover seeds. The spacing between probe sites is set from the covering radius of the lattice, which is what allows the method to state in advance the size below which a cluster may escape detection. The second phase, an expansion refinement activated only on probes that return an occupied voxel, verifies every edge, so the components returned are true connected components. BLS’s versatility allows for selection of different Bravais lattice unit cells to match the target structure; for amorphous, non-crystalline shapes, BLS can default to a simple face-centred cubic unit cell, where the expected minimum cluster size is the only parameter that needs to be set. The current BLS implementation has been developed as a post-processing tool for molecular dynamics trajectories, and was tested for searching water ice clusters of different morphologies. BLS returns component counts and maximum cluster sizes identical to exhaustive-labeller algorithms, with 100% recall; it runs at about 0.94 times the cost of depth-first search, and at 0.84 to 0.90 times the cost of the fastest other labeller in our benchmark set. This algorithm, although implemented by us for molecular dynamics applications, could be of interest in other domain areas where searching for high-density elements in 3D space is relevant.

## 1 Introduction

Identifying connected high-density regions, in three-dimensional discrete spaces is a basic operation in crystallography, molecular dynamics, medical imaging and materials science. Given a binary voxel grid 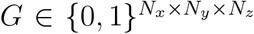, the task is to partition occupied voxels (*G*_*i,j,k*_ = 1) into disjoint connected components under a specified neighbourhood relation (clusters). This is the connected component labeling (CCL) problem, and it has well-established exact solutions [1, 2]. They share one structural commitment: every voxel, or at minimum every occupied voxel, is visited. In nucleation studies *in-silico*, for instance, the crystalline phase often occupies a smaller fraction of the simulation volume [15]. That is the opening a sampling-based method can exploit, and it is what motivates the approach taken here.

### 1.1 Algorithm Families Considered

BLS is compared against a set of algorithms drawn from four families. Two of these families do not solve the CCL problem at all. However we include them because BLS performs first a search followed by clustering. Thus it is informative to see how methods designed for one of those two different problems behave when handed the whole job. We do not aim to perform a full survey of benchmarks; for those interested, the literature is reviewed elsewhere [1]. Below we list the algorithms taken into account, grouped by family.

#### Exact connected-component labeling

Depth-first search (DFS) visits every occupied voxel once, at *O* (*N*_occ_) cost [3]. We include DFS, a union-find labeler of the kind provided by the `cc3d` library [4], GCBD [5] and run-length-encoded RLE-CCL [6]. These solve exactly the problem BLS solves and define the correct answer: any method claiming exactness must reproduce their output component for component. We have taken this family of algorithms as the primary comparison and is used to validates correctness aswell.

#### Seed-and-grow

This family is close to BLS in the implementation itself: VCCS [7] places seeds on a uniform grid and grows regions outward, and shares the two-phase shape of BLS. It is therefore the most informative comparison in the paper, because the two methods differ in one ingredient only: VCCS chooses its seed spacing independently of the objects being sought and offers no statement about what it can miss, whereas BLS fixes the spacing from a covering radius, provided by the user. See Section 2.1.

#### Density-based

DBSCAN [8] and HDBSCAN [9] treat clusters as dense regions separated by sparse ones, controlled by a neighbourhood radius and a minimum point count. They do not check that a connected path of occupied voxels joins two points placed in the same cluster; and their parameters have no direct correspondence to crystallographic length scales. They therefore answer a related but distinct question. Including them measures what connection inference costs relative to verification, particularly as density rises.

#### Partitioning

This family of algorithms deals with a different problem, reported here for completeness. *k*-means [10] and agglomerative hierarchical clustering divide data into *k* groups by optimizing a geometric objective. *k*-means requires *k* in advance and agglomerative clustering a distance threshold, so both are handed information that the connectivity methods must discover, and they carry no notion of voxel adjacency. They do not solve the CCL problem. We ran them because they are what a practitioner reaches for by default when asked to cluster points in space, and their failure modes are worth documenting; their results appear in the Supplementary Information.

**Table 1:**
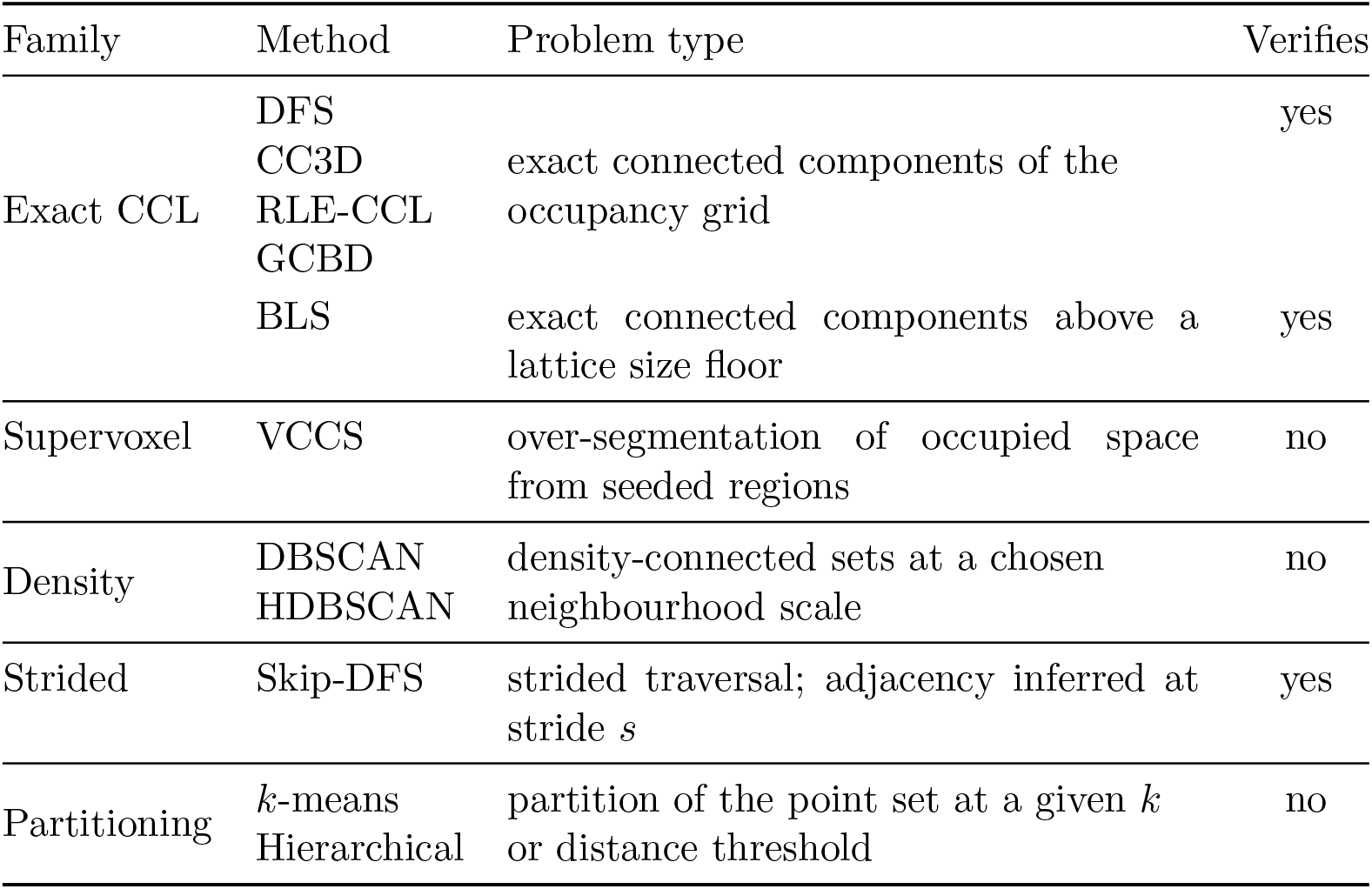
The comparison set, grouped by the problem each family solves. *Verifies* column records whether a connected path of occupied voxels is checked. Partitioning methods are reported in the Supplementary Information only.

### 1.2 Bravais Lattices

A Bravais lattice Λ is the infinite set of points reached by integer steps along three basis vectors **a**_1_, **a**_2_, **a**_3_:

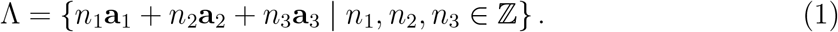

The 14 three-dimensional Bravais lattices are classified by crystal system and centering [11]: primitive (P), one lattice point per conventional cell; body-centred (I), contains two points; face-centred (F), four. Two length scales describe the same lattice, and their ratio depends on the centering: the conventional cell parameter *a*, the edge of the cube containing one conventional cell, and the nearest-neighbour distance *d*_NN_ between adjacent sites.

### 1.3 Connected Component Labeling

Given a binary grid *G* = {0, 1}^*M×M×M*^ , where *G*_*i,j,k*_ = 1 denotes an occupied voxel, and a neighbourhood relation *N*, CCL assigns a unique label to each maximal connected subset of occupied voxels. We use 6-connectivity throughout:

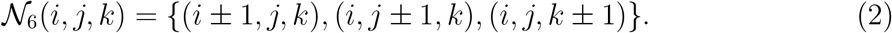

Conventional CCL locates its seeds by rastering the grid, at cost *O* (*M* ^3^) regardless of how little of the grid is occupied.

## 2 Proposed Method

BLS separates the two tasks the raster scan: finding one point inside each structure, and recovering the structure around it. Seeds are discovered by evaluating a sparse set of probe sites placed on a Bravais lattice (Section 2.1); each occupied probe is then expanded into a component by depth-first traversal on the full grid (Section 2.2). Only the seeding phase is subsampled, this allows BLS for a potential faster execution.

### 2.1 Geometry-Guided Sparse Probing

Probe sites are placed at Bravais lattice positions scaled so that *d*_NN_ matches the characteristic diameter of the structures sought. The covering radius of the lattice, that is the largest distance from any point in space to the nearest probe site, bounds the size of a structure that can escape detection. Expressed in units of *d*_NN_, the covering radii of the three cubic centerings are

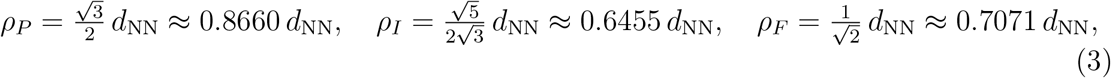

so for the face-centred cubic lattice any region containing a ball of diameter 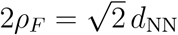 contains at least one probe site. This is a sufficient condition on a continuous region.

Comparing centerings at a fixed *d*_NN_ gives them different probe budgets. The quantity that compares them at equal cost is the covering radius normalized by the probe density *v*, 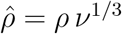, which for the same three centerings is

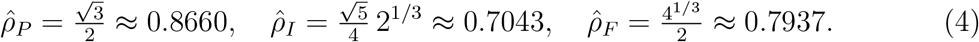

The body-centred lattice is the thinnest lattice covering of three-space [12, 13], which is the ordering 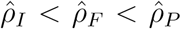. Section 4.7 tests whether that ordering and its magnitude appear in detection.

BLS derives the spacing from a covering radius. The price is a size floor that is stated and computable. For nucleation analysis, where sub-critical nuclei are discarded anyway, that floor is the physical radii of the expected minimum cluster size.

#### 2.1.1 FCC Lattice Formulation

For the face-centred cubic lattice the basis vectors are

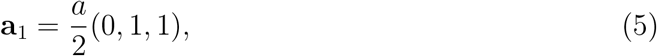

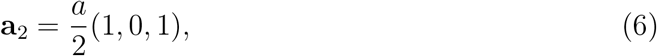

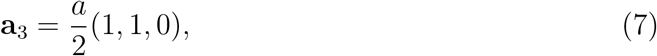

with 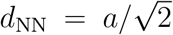. The user supplies the target diameter through *d*_NN_ , which fixes 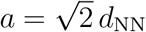.

#### 2.1.2 Probe Count

An *F* -centred conventional cell of edge *a*, expressed in voxels, carries four lattice points, so the probe density is 4*/a*^3^ and the *M* ^3^ voxels of the grid receive

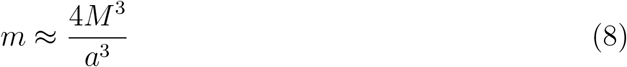

probe sites, each evaluated in *O* (1). These produce the seed set *S* = {(*i, j, k*) ∈ Λ | *G*_*i,j,k*_ = 1}, replacing the *O*(*M* ^3^) raster scan of Section 1.3.

### 2.2 High-Resolution Refinement

Each seed in *S* is expanded by depth-first traversal under *N*_6_ on the full-resolution grid, marking visited voxels so that no voxel is claimed twice. Expansion terminates when the frontier is empty, at which point the traversal has enumerated one complete connected component. Every edge is tested, so the components returned are true connected components and BLS agrees with an exhaustive labeler on every component it reaches.

The traversal admits a stride parameter *s*, which replaces *N*_6_ with *N*_*s*_(*i, j, k*) = {(*i* ± *s, j, k*), (*i, j*±*s, k*), (*i, j, k*±*s*)} and infers connectivity from proximity rather than verifying it. Every BLS result reported in this paper uses *s* = 1, for which _*s*_ = _6_ and nothing is skipped. In other words, when *s* = 1 the expansion goes with a standard DFS algorithm. The *s >* 1 case is a separate algorithm, Skip-DFS, measured in Section 4.6.

### 2.3 Parameters

A BLS run is specified by the following settings. grid spacing is the voxel edge in Å and fixes the resolution at which occupancy is discretized, so it bounds both the cost and the smallest separation at which two clusters can still be told apart. radii are the van der Waals radii used to mark the voxels each atom occupies. cutoff is the fraction of a voxel that must be covered before it counts as occupied, and occupancy selects whether any atom or every atom of the group is required. lattice and centering select the Bravais lattice and centering of the probe set (Section 2.1). connectivity is the neighbourhood relation used by the refinement, *N*_6_ throughout, and skip is the traversal stride *s* of Section 2.2, *s* = 1 throughout. box defines the cell; `AUTO` fits a bounding box to the atoms with a padding of twice the grid spacing.

The probe spacing is set by dnn and ALPHA. dnn gives *d*_NN_ directly in Å; Setting `DNN = 0` derives it instead as

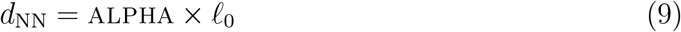

so that ALPHA is a dimensionless scale factor on a fixed reference length *ℓ*_0_. It is therefore the single knob that sets how coarse the probe lattice is relative to the structure being searched: ALPHA = 1 places probes one reference length apart, larger values spread them further. Since the number of probes falls as 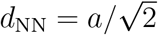, increasing ALPHA lowers the seeding cost, but the covering radius grows with it, so above some system-dependent ALPHA^*\**^ the coarsest probes no longer land inside the smallest clusters and recall drops. The operating point is the largest ALPHA at which recall is still exact; because the probe spacing enters in voxels, ALPHA^*\**^ is defined only jointly with grid spacing and must always be quoted with it.

### 2.4 Time Complexity

- **Probing**: with *n* = *a*^3^*/*4 voxels per probe cell (set by the target cluster size), the number of probes is *m* = *M* ^3^*/n*, giving a probing cost of *O*(*M* ^3^*/n*).
- **Refinement**: *O*(*αM* ^3^) at unit stride, where *α* ∈ [0, 1] is the occupied fraction of the grid.
- **Total**: *O*(*M* ^3^*/n*) for sparse grids (*α* ≪ 1), degrading toward exhaustive behaviour as *α* → 1.
- **Probing**: with *n* = *a*^3^*/*4 voxels per probe cell (set by the target cluster size), the number of probes is *m* = *M* ^3^*/n*, giving a probing cost of *O*(*M* ^3^*/n*).
- **Refinement**: *O*(*αM* ^3^) at unit stride, where *α* ∈ [0, 1] is the occupied fraction of the grid.
- **Total**: *O*(*M* ^3^*/n* + *αM* ^3^), degrading toward exhaustive behaviour *O*(*M* ^3^) as *α* → 1.

## 3 Experimental Setup

### 3.1 Implementation and Software

BLS and all comparison algorithms were implemented in C++ without external numerical libraries. The benchmark harness, trajectory readers, comparison algorithm implementations, and analysis pipeline were developed with the assistance of Claude Code (Anthropic), an agentic coding assistant. All algorithmic design decisions, parameter choices, experimental protocols, result interpretation, and scientific claims are the author’s; the assistant was used for code generation, refactoring, and data processing under direct supervision. All algorithms operate on raw voxel adjacency, and every result below is to be read on that basis. Every implementation was verified against unit tests covering lattice geometry, connectivity modes, determinism, and cross-algorithm output agreement, with every geometric test exercising a non-orthogonal, non-symmetric triclinic basis in addition to the cubic one. Source code, structures used for in-silico experimetns, configuration files, and results are available at https://github.com/franciscocarrascoza/BLS.

Development and some measurements were taken on an Intel Core i7-13700 under a Release build (`-O3 -DNDEBUG`), single-threaded, with the process pinned to the performance-core set. Replication of results were verified on a dual-socket Intel Xeon Platinum 8268 (Cascade Lake) system, 2 sockets × 24 cores, 48 physical cores in total with one thread per core (hyperthreading disabled), running at a base clock of 2.90 GHz and boosting to 3.90 GHz.

The settings used for most of the BLS runs are reported below in the format of the input file used

~~~
BLS …
   GROUP ATOMS=all
   BOX AUTO
   GRID SPACING 4.0
   CONNECTIVITY 6
   SKIP 1
   ALPHA 2.45
   DNN 0
   RADII 1.5, 1.2
   CUTOFF 0.25
   OCCUPANCY ANY
   LATTICE cubic
   CENTERING F
…BLS
~~~

except where a parameter is the one being swept. Section 4.1 reports how grid spacing, ALPHA, lattice and centering were fixed at these values; the remaining settings are carried over from E0. The comparison algorithms take GRID_SPACING, RADII, CUTOFF and CONNECTIVITY from the same deck, so all methods label the same grid.

### 3.2 Benchmarking Fairness

Each published method is benchmarked in the form its own description specifies, including the data structures that description assumes. For CC3D that is the two-scan SAUF procedure [2] with a rolling two-plane label buffer; for RLE-CCL it is a union-find over runs [6]; for VCCS it is adaptive seeding with seed pruning [7]. These structures are the mechanism by which each method achieves the complexity its authors claim. Stripping them out yields a different algorithm. The rest of the benchmarked algorithms, including BLS, where implemented with its basic text book implementation. Meaning optimized resources and libraries were avoided. GCBD is reported in a single implementation, since no distinct tuned form is defined for it. Every method labels the same grid, produced by the same voxelisation from the same structure file, and all timings are wall-clock over the labelling call alone.

Two implementation notes bear on interpretation. The `cc3d` reference library is distributed under LGPL-3.0-or-later and is header-only, so any use would be static incorporation and incompatible with the MIT licence of this work; CC3D was therefore reimplemented from the published description rather than linked. VCCS was ported rather than linked, because the reference implementation pulls in four large dependencies for one algorithm, and its colour and normal feature terms were omitted: a binary occupancy grid has no colour, and a surface normal is undefined for an occupancy indicator, so only the spatial term carries information.

### 3.3 Test Systems

Water ice was chosen as the validation material for two reasons. Its polymorphs are well known from experimentally resolved crystal structures with accurately known bulk densities, which turns the systems into a control on the measurement itself rather than only a substrate for it. And cluster detection in ice is a real application the method can tackle, since nucleation analysis asks exactly the question BLS answers, how many separate supercritical nuclei are present and how large the largest is. Table 2 reports the measured densities. The ice controls agree with their bulk references to within 2% for I_c_ and VII, which is what licenses the use of the same measurement on the disordered system. The disordered system is packed to 0.020 83 mol Å^*−*3^ , 67% of the low-density amorphous reference, and is reported throughout by that measured density as low-density disordered water. It probes the detection limit at low occupancy, and the geometry is what the experiments use.

**Table 2:**
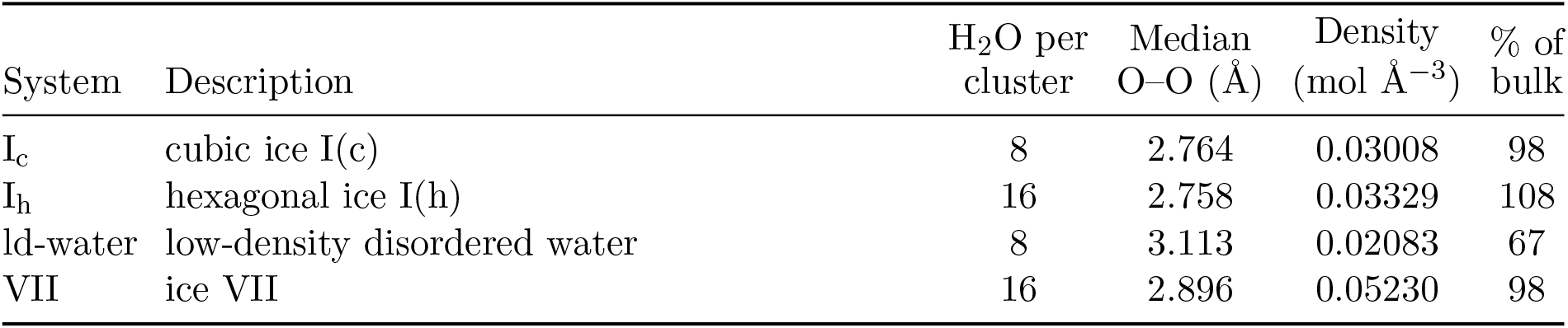
Test systems and their measured densities. The bulk reference is 0.0307 mol Å^*−*3^ for the ice I polymorphs, 0.0313 mol Å^*−*3^ for low-density amorphous ice and 0.0535 mol Å^*−*3^ for ice VII. The ice controls validate the measurement itself. The disordered system sits at 67% of the low-density amorphous reference and is reported as low-density disordered water. All six E2 containers are packed from that same motif, so E2 varies the container and holds the material fixed.

The disordered system measures 0.020 83 mol Å^*−*3^ , 67% of the low-density amorphous reference, and is reported throughout by that measured density as low-density disordered water. It probes the detection limit at low occupancy, and the geometry is what the experiments use.

### 3.4 Experiments

The test systems were used to create seven different experiments, where E0 and E6 were meant to search the optimal parameters of BLS. The rest of the experiments tested every different aspect of the problem BLS aims to solve: search and cluster.

#### E0. Parameter sensitivity and deck selection

Two parts. A sensitivity sweep of GRID_SPACING, SKIP, CUTOFF, ALPHA, LATTICE and CENTERING in three phases, on three systems (Ice I_c_, Ice I_h_, low-density disordered water). Then a joint search over grid spacing and ALPHA on all four systems for the setting of least runtime that holds recall at exactly 1.0 on every one of them, worst case over eight lattice-origin offsets (*k, k, k*) voxels, *k* = 0 … 7: 2560 runs on a coarse ALPHA grid and 800 on a refinement, at five replicates, followed by a repetition of the search at each of five grid spacings. The component count present is re-measured by DFS at each grid spacing, since it is grid-dependent.

#### E1. Ice polymorphs

All algorithms on four ice systems (I_c_, I_h_, low-density disordered water, VII), 1000 packed clusters each. This experiment examines impact of the target shape on detection accuracy using a given lattice.

#### E2. Packed-cloud shape

Six container shapes (cube, cylinder, sphere, triclinic, dodecahedron, octahedron), 100 clusters each. None of the six carries periodic boundary conditions defined (i.e., CRYST1 records), so all six fall through to a diagonal autobox: the box matrix is the same in every case and only the shape of the packed atom cloud varies. E2 therefore tests invariance to the shape of the packed cloud inside an orthorhombic cell, and is reported in the Supplementary Information on that basis.

#### E3. Scaling

System size swept from *n* = 1 to *n* = 5000 expected clusters, with the volume growing so that density stays fixed.

#### E4. Density transition

Cluster count swept from *n* = 1 to *n* = 750 in a fixed box, crossing the percolation threshold.

#### E5. Gap resolution

Two-cluster and 20-cluster systems with inter-cluster gaps from 1 to 20 Å.

#### E6. Probe-budget-normalized lattice comparison

Cluster diameter swept downward through *d*_NN_ at equalized probe count for centerings P, I and F, recording the diameter at which recall departs from 1.0. Unlike E0, which held *d*_NN_ fixed and therefore gave the three centerings different probe budgets, E6 equalizes cost and is the configuration in which Equation (4) can be tested. A primary arm equalizes the budget through ALPHA on the four E1 systems; an independent control arm sweeps the diameter of 216 random-placed clusters in a fixed box. Both arms sweep eight lattice-origin offsets and report the worst case.

## 4 Results

Every configuration reported below was executed as five independent replicate processes. Detection is deterministic: repeating a configuration on the same input reproduces the detected counts and the maximum cluster sizes bit for bit, verified across 672 replicate groups during the deck search with no exceptions, so results resting on discrete quantities are reported without a run-to-run uncertainty. Timings are the median over the five replicates and carry the replicate spread beside them. The comparison of Section 4.3 is 54 algorithm–system configurations, 270 processes, with no failures. Runtime and recall are coupled through grid spacing, so every timing below states the spacing at which it was measured and transfers to another spacing only on re-measurement.

### 4.1 Sampling Geometry and the Working Deck

Of the six BLS parameters measured in E0, two govern behaviour and four are nearly inert over the tested ranges (Table 3, Figure 2a). When testing lattice, the best-case recall drops by 0.210, from 0.990 for cubic to 0.780 for triclinic, and by 0.120 across centering, from 0.990 for *F* to 0.870 for *P* . The remaining four parameters change recall by at most 0.009. Within that group ALPHA and GRID_SPACING carry a mild accuracy–speed trade-off, recall falling by 0.007–0.009 as either is coarsened; cutoff changes recall by 0.006; and skip is flat at 0.992 across all five tested values, the expected signature of a parameter that takes effect only above unit stride. Recall is reported as the minimum across the three systems: at the finest grid spacings the disordered system over-segments and detects roughly twice the expected number of clusters, which inflates a mean without indicating better detection. The sampling geometry therefore has to be chosen before anything else, and it is chosen once. Figure 2(b) resolves the centering at the working deck, where all four systems and eight lattice-origin offsets are swept. Cubic-*P* misses two of the 988 components on ice I_c_ at its worst offset. Cubic-*I* and cubic-*F* both reach recall 1.000 on all four systems. The face-centred default is admissible; the measurement does not single it out, since the body-centred lattice performs identically here and was not timed.

**Table 3:**
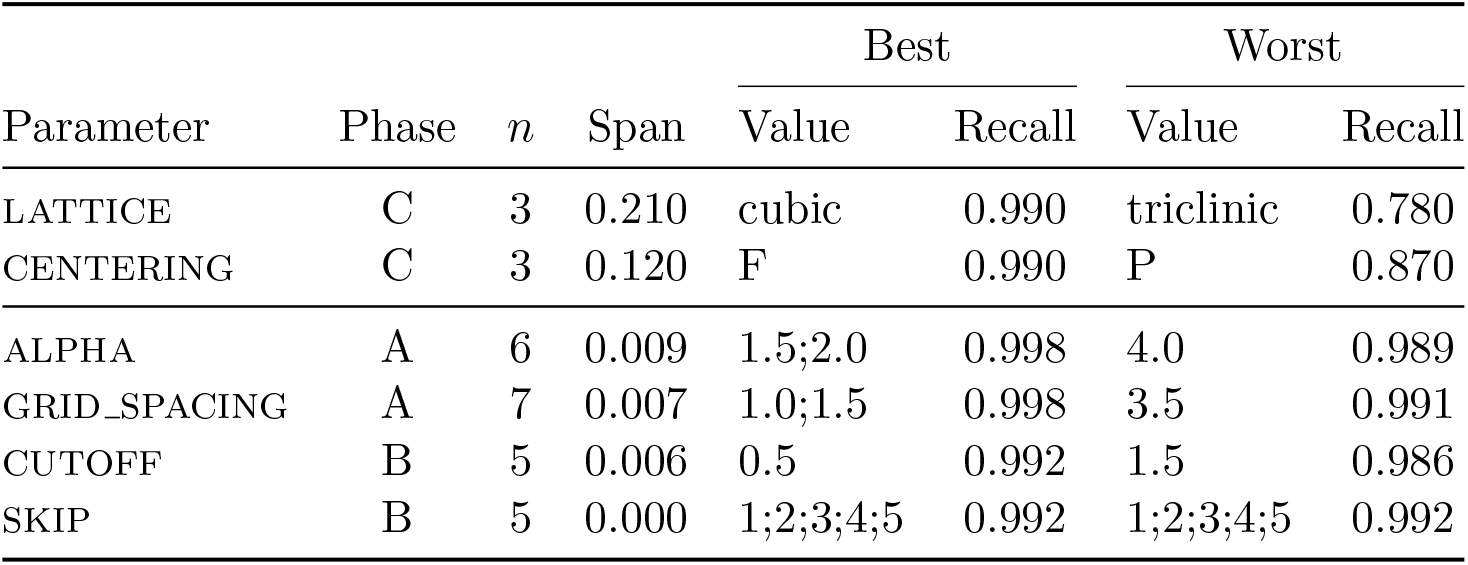
BLS parameter sensitivity (E0), measured at a 3.5 Å grid spacing. Best-case recall at a value *v* is the maximum, over every tested combination holding the parameter at *v*, of that combination’s minimum recall across the three systems; *span* is the range of that quantity over the tested values. The minimum rather than the mean is used because the mean exceeds unity wherever a setting over-segments. The two parameters above the rule select the sampling geometry; the four below control rasterisation and traversal. *Phase* identifies which of the three E0 experimental phases (A, B, C) tested that parameter.

**Figure 1:**
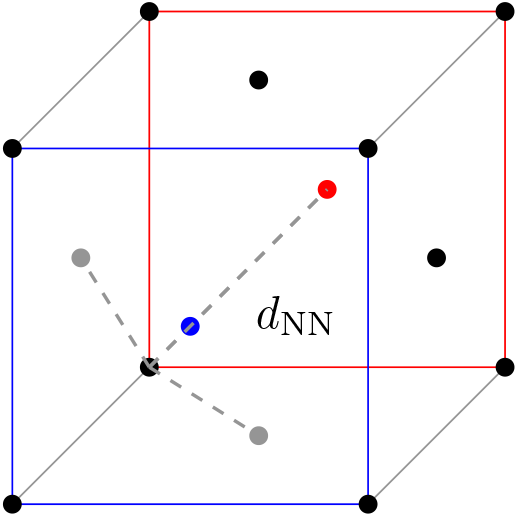
Face-centred cubic (FCC) sampling cell. Dots are probe sites (8 corner + 6 face-centred). The dashed line marks the distance from the origin to its nearest neighbours, 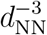.

**Figure 2:**
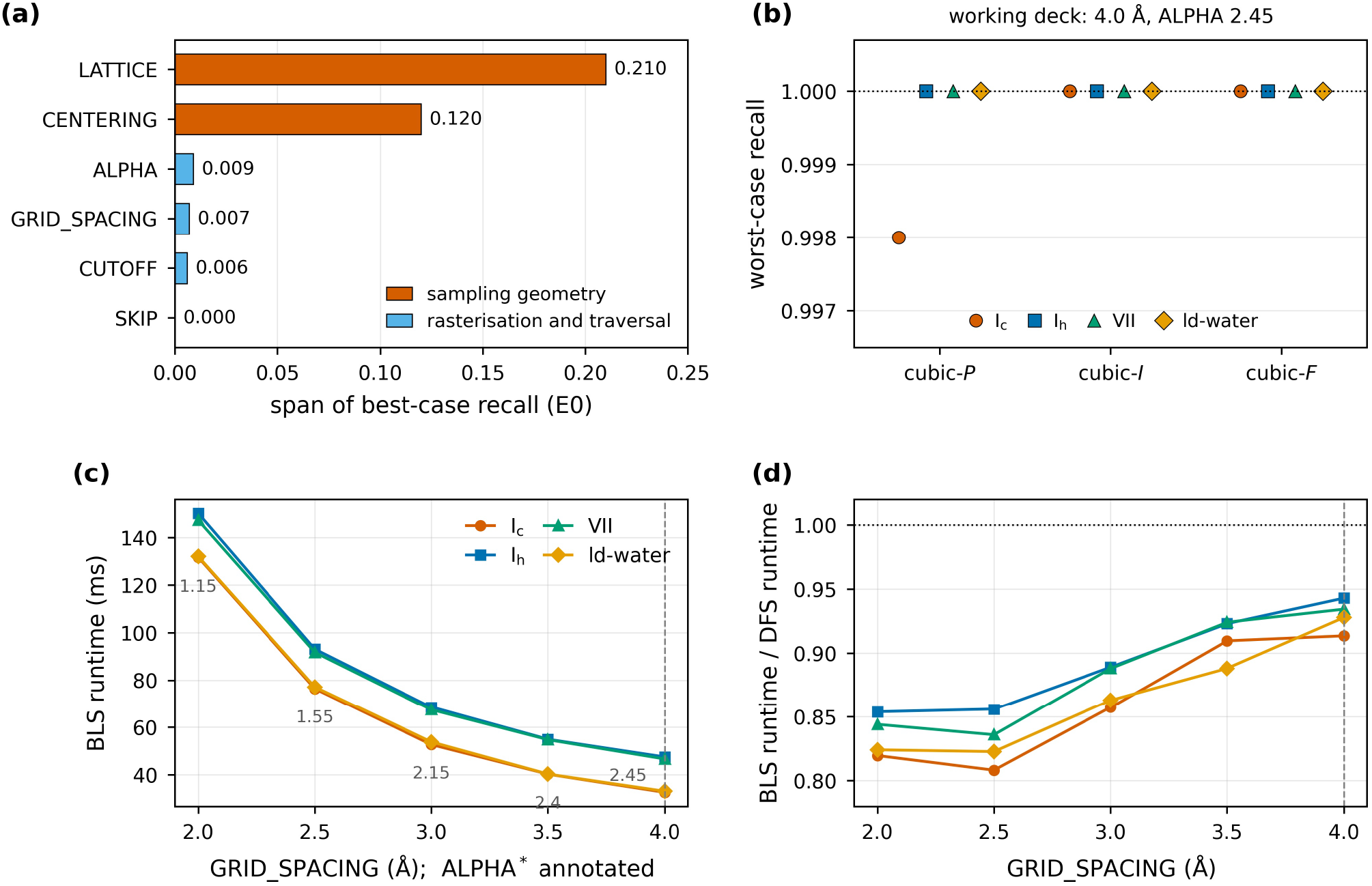
BLS parameters and the working deck. (a) Span of best-case recall over the tested values of each E0 parameter, at 3.5 Å. The two sampling-geometry parameters exceed the four rasterisation and traversal parameters by more than an order of magnitude. (b) Worst-case recall over eight lattice-origin offsets for the three cubic centerings at the working deck; markers are offset horizontally so that coincident values remain visible. (c) BLS runtime against grid spacing, each spacing at its own ALPHA^*\**^, the coarsest probe lattice holding recall at 1.0000 on all four systems; the annotated values are ALPHA^*\**^ and the dashed rule marks the working deck. (d) The same runs as a ratio to DFS measured on the same grid.

The remaining two parameters were fixed by a joint search under a single constraint: recall exactly 1.000 on all four systems, worst case over eight origin offsets, with runtime minimised subject to it. The boundary in ALPHA moves with the grid, from ALPHA^*\**^ = 1.15 at 2.0 Å to 2.45 at 4.0 Å (Figure 2c). Ice I_c_ and the disordered system are the binding cases at every spacing; ice I_h_ and ice VII alone tolerate ALPHA up to 3.2–3.3. All five spacings reach recall 1.0000 at their own boundary, so the choice among them is runtime alone, and BLS runtime falls monotonically as the grid coarsens on every system individually. The working deck is therefore

~~~
GRID_SPACING 4.0, ALPHA 2.45, LATTICE cubic,
CENTERING F, CUTOFF 0.25, SKIP 1,
~~~

uniform across all four systems, and every number in Sections 4.2 and 4.3 is measured there.

Figure 2(d) records the cost of that choice. BLS runs at 0.808–0.856 times DFS at grid spacing of 2.5 Å and at 0.917–0.958 times DFS at 4.0 Å. Therefore the relative margin narrows as the grid coarsens while the absolute runtime falls by a factor of two and a half over the same range. The working deck is the fastest deck in absolute terms and the one at which the ratio to DFS is closest to unity. Both statements hold at recall 1.000.

At the working deck the probe lattice has *d*_NN_ = 1.47 voxels, so Equation (8) gives 4*/a*^3^ = 0.445 probe sites per voxel, a 2.25-fold reduction in the number of sites examined during seeding relative to a raster scan of the same grid. The measurement agrees: on ice I_c_, 8291 probe sites fall on occupied voxels against 18 252 occupied voxels present, a ratio of 0.454. The runtime saving that this buys is reported in Section 4.3 and is smaller than the reduction in site count.

### 4.2 Accuracy Against the Comparison Set

On every component it reaches, BLS agrees with an exhaustive labeler exactly. Over a matrix of 192 cells, four systems by six lattice–centering combinations by eight lattice-origin offsets, no component was truncated, split or merged in any cell, and the number of clusters BLS reports equals its seeded-component count in all 192. The maximum cluster size matches DFS in 191 of the 192 cells; the exception is the hexagonal-*P* lattice on ice VII, where the largest component was never seeded, so the largest BLS returns is the next one down. That is a seeding outcome, and it does not occur at the working deck.

Table 4 gives the exact-connectivity tier at the working deck. BLS, DFS, CC3D, RLE-CCL and GCBD return identical component counts and identical maximum cluster sizes on all four systems, and recall is exactly 1.0000 for each of them. Recall divides by the component count DFS measures on the same grid, which is 989, 991, 990 and 994 against a nominal packing request of 1000 clusters; random packing overlaps and merge a handful of clusters, so the requested figure was never the ground truth. Coarsening the grid from 3.5 Å to 4.0 Å merges a few more components, which moves the present count by one to four per system, so counts measured at the two spacings are not comparable cluster for cluster.

**Table 4:** Exact-connectivity tier on the four E1 systems at the working deck (4.0 Å grid spacing, ALPHA = 2.45, cubic-*F*). Times are the median of five replicate processes and spread is (max − min)*/*median over those replicates. Detected counts are given against the component count present, measured by DFS on the same grid, which is what recall divides by. RSS is peak resident set size in megabytes; Max. cl. is the size of the largest detected cluster.

| System | Algorithm | Detected / present | Recall | Time (ms) | Spread (%) | $t/t_{\text{DFS}}$ | RSS (MB) | Max. cl. |
| --- | --- | --- | --- | --- | --- | --- | --- | --- |
| $I_c$ | BLS | 989 / 989 | 1.0000 | 28.2 | 5.6 | 0.944 | 36 | 41 |
|  | DFS | 989 / 989 | 1.0000 | 29.8 | 8.8 | 1.000 | 35 | 41 |
|  | CC3D | 989 / 989 | 1.0000 | 33.7 | 4.8 | 1.130 | 36 | 41 |
|  | RLE-CCL | 989 / 989 | 1.0000 | 34.7 | 4.1 | 1.162 | 35 | 41 |
|  | GCBD | 989 / 989 | 1.0000 | 102.3 | 2.0 | 3.427 | 218 | 41 |
| $I_h$ | BLS | 991 / 991 | 1.0000 | 43.5 | 4.4 | 0.958 | 36 | 57 |
|  | DFS | 991 / 991 | 1.0000 | 45.4 | 3.0 | 1.000 | 36 | 57 |
|  | CC3D | 991 / 991 | 1.0000 | 48.1 | 3.6 | 1.060 | 36 | 57 |
|  | RLE-CCL | 991 / 991 | 1.0000 | 48.3 | 3.9 | 1.062 | 36 | 57 |
|  | GCBD | 991 / 991 | 1.0000 | 117.7 | 1.1 | 2.591 | 219 | 57 |
| VII | BLS | 994 / 994 | 1.0000 | 42.4 | 2.9 | 0.949 | 36 | 52 |
|  | DFS | 994 / 994 | 1.0000 | 44.7 | 3.8 | 1.000 | 36 | 52 |
|  | CC3D | 994 / 994 | 1.0000 | 47.9 | 2.5 | 1.070 | 36 | 52 |
|  | RLE-CCL | 994 / 994 | 1.0000 | 48.0 | 3.6 | 1.074 | 36 | 52 |
|  | GCBD | 994 / 994 | 1.0000 | 116.7 | 1.1 | 2.610 | 218 | 52 |
| ld-water | BLS | 990 / 990 | 1.0000 | 28.1 | 7.2 | 0.917 | 35 | 59 |
|  | DFS | 990 / 990 | 1.0000 | 30.7 | 5.4 | 1.000 | 35 | 59 |
|  | CC3D | 990 / 990 | 1.0000 | 33.1 | 4.4 | 1.081 | 36 | 59 |
|  | RLE-CCL | 990 / 990 | 1.0000 | 34.4 | 3.9 | 1.122 | 35 | 59 |
|  | GCBD | 990 / 990 | 1.0000 | 103.4 | 2.5 | 3.370 | 218 | 59 |

Figure 3(b) places the rest of the comparison set on the same axes. *k*-means returns 0.997–1.004 of the present count, with *k* supplied to it. Skip-DFS at unit stride returns 0.987–0.995, DBSCAN 0.971–0.984 and agglomerative clustering 0.961–0.973. VCCS returns 4.49–5.45 times the present count and HDBSCAN returns a single component on both systems where it completed, a recall of 0.001. Only the five methods of Table 4 land on the correct count.

**Figure 3:**
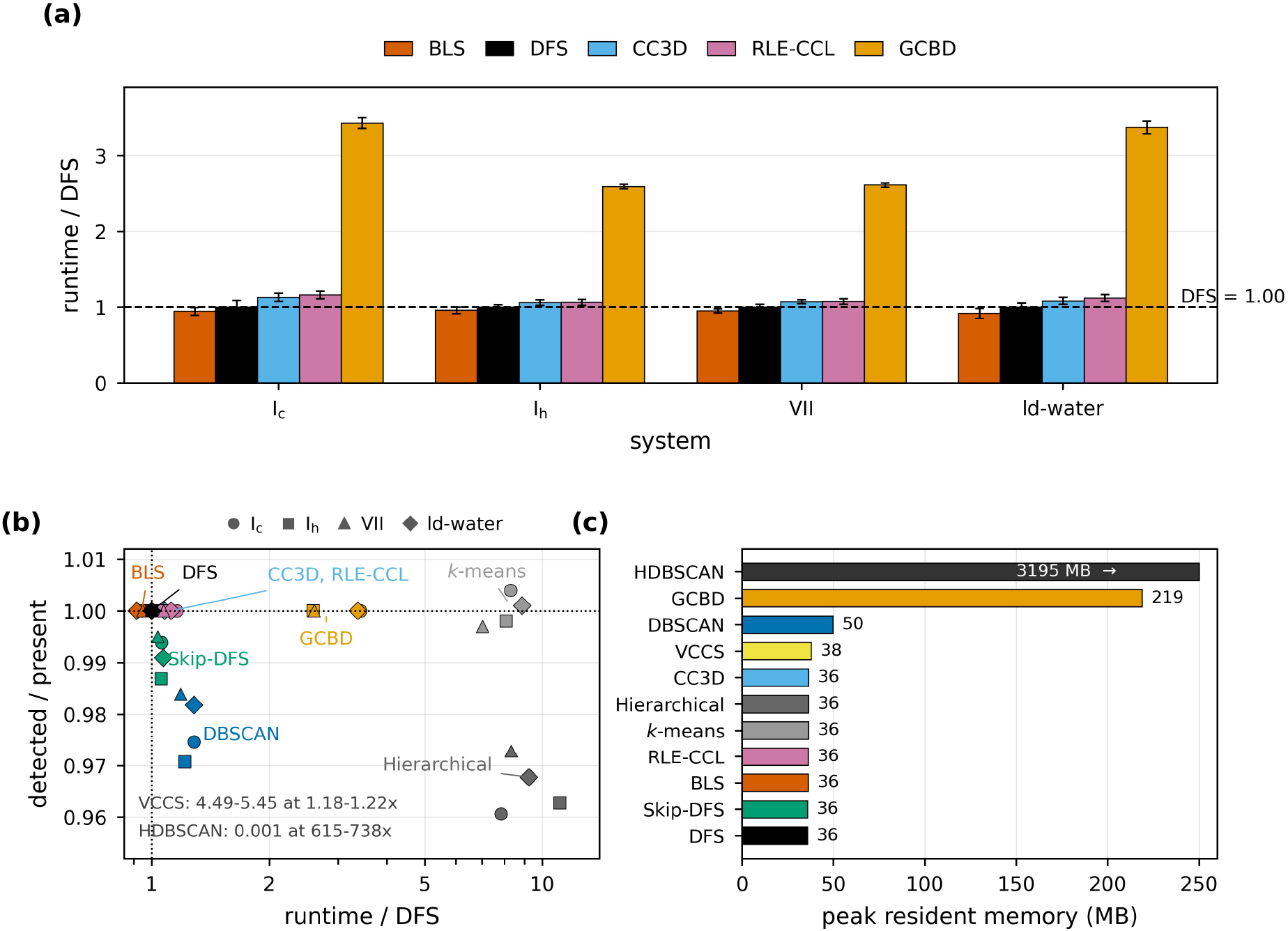
BLS against the comparison set at the working deck (4.0 Å grid spacing). (a) Runtime normalized to DFS on each system; error bars are the replicate spread over five replicate processes. (b) Detected component count divided by the count present, against runtime ratio, for every method reported in the main text. Marker shape gives the system. VCCS and HDBSCAN fall outside the plotted window and their values are stated in the panel. (c) Peak resident memory, the maximum over the four systems; the HDBSCAN bar is clipped and its value annotated.

The VCCS figure is the designed behaviour of a supervoxel method [7] and not a detection failure. Its output count is set by the seed resolution *R*_seed_ and tracks the number of seeds placed at a ratio of 1.0000 on all four systems; over the seed-resolution sweep the over-segmentation ratio falls monotonically from 7.85 at *R*_seed_ = 2 to 1.00 at *R*_seed_ = 12. Both VCCS and BLS place seeds, grow regions, and admit a spacing that controls how many objects they return. The full sweep is given in the Supplementary Information.

### 4.3 Speed and Memory

At the working deck BLS runs at 0.917–0.958 times DFS on the four systems and is the fastest method in the comparison (Figure 3a, Table 4). CC3D and RLE-CCL cost 1.06–1.13 and 1.06–1.16 times DFS, Skip-DFS 1.04–1.07, VCCS 1.18–1.22, DBSCAN 1.19–1.28 and GCBD 2.59–3.43. The partitioning methods cost 7.0–11.1 times DFS and HDBSCAN 615–738 times.

The margin over DFS is 4.2% to 8.3% against a replicate spread of 2.9% to 8.9% at this deck. The sign of the margin held on all four systems here and in every earlier measurement at every grid spacing tested (Figure 2d), so the direction is safe; the magnitude is at the resolution limit on ice I_h_ and ice VII and we quote it to two significant figures nowhere.

Peak resident memory separates into two groups (Figure 3c). BLS, DFS, CC3D, RLE-CCL, Skip-DFS, *k*-means and agglomerative clustering all sit between 35 and 36 MB at E1 size, which is the occupancy grid itself. DBSCAN reaches 49 MB, VCCS 38 MB, and GCBD 218–219 MB, six times the grid, from the block bookkeeping its decomposition maintains. HDBSCAN reaches 2577 MB on ice I_c_ and 3195 MB on the disordered system, and did not complete on the two denser systems; its mutual-reachability edge list grows as the square of the occupied voxel count.

BLS and DFS share a traversal proportional to occupancy, and the two sit in one performance tier while GCBD sits in another. What separates BLS from DFS within that tier is the seed scan, and Section 4.1 measured what it removes: 55% of the site visits, for a 4–8% saving in total runtime. The seeding phase is therefore a minority of the cost at these occupancies, and the probe sites are visited in lattice order while a raster scan reads the grid in memory order.

### 4.4 The Size Floor

BLS answers a parameterized query. It find the components whose size exceeds a floor set by *d*_NN_, which is a user input. Components below that floor lie outside the question asked, in the same sense that the neighbourhood radius of DBSCAN and the seed resolution of VCCS bound what those methods return. A recall figure compares BLS against a method answering an unparameterized question, and should be read with that asymmetry in mind.

Figure 4(a) gives the measured floor at *d*_NN_ = 2.4 voxels, pooled over the 192-cell matrix; a component size counts as missed if it was missed in any cell at any origin offset, so the curve is a worst case. The miss rate falls monotonically with component size across the well-sampled range and reaches zero above 60 voxels. The largest component ever missed is 59 voxels, which is a floor of 60 pooled over all six lattices, including hexagonal-*P* . Restricted to the cubic-*F* default the largest missed is 28, a floor of 29.

**Figure 4:**
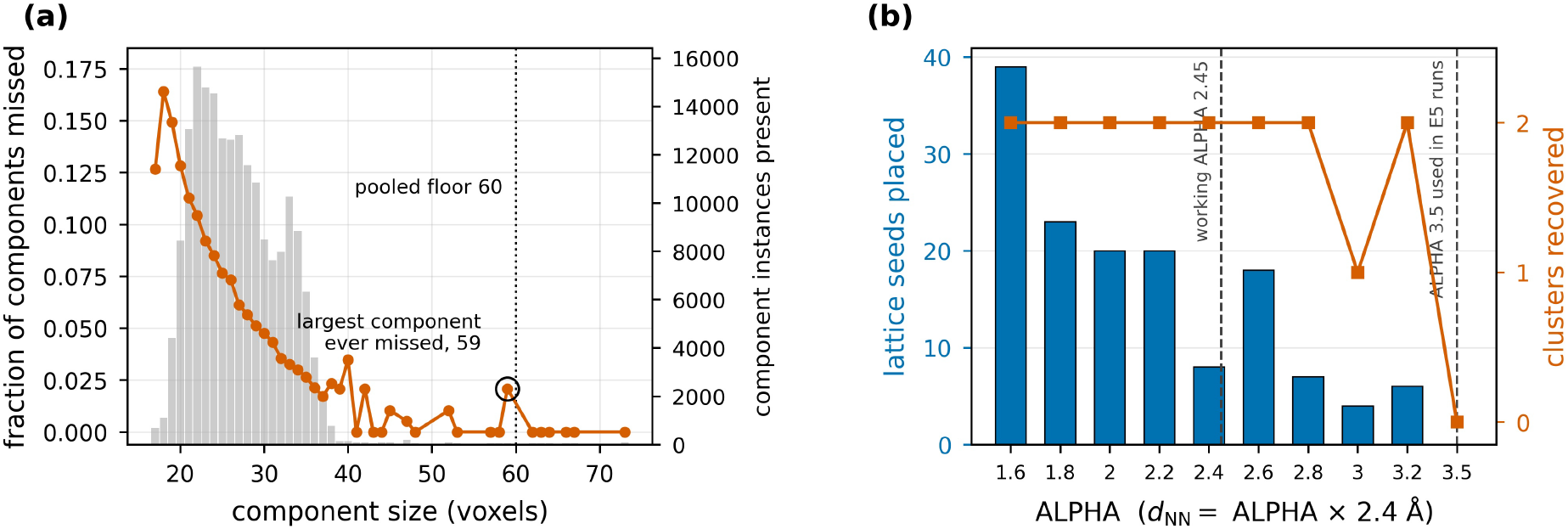
The BLS size floor. (a) Fraction of components missed against component size, pooled over the 192-cell matrix of four systems, six lattice–centering combinations and eight lattice-origin offsets, at *d*_NN_ = 2.4 voxels; a size counts as missed if it was missed in any cell. Grey bars give the number of component instances of each size present, which is the support the rate is measured on. The dotted rule is the 60-voxel floor pooled over all six lattices and the circled point is the largest component ever missed. (b) The same floor on the two-cluster E5 system at a 20 Å gap and a 3.5 Å grid spacing, sweeping ALPHA and therefore *d*_NN_. Both clusters are recovered at ALPHA ≤ 3.2, which includes the working value.

The floor scales with *d*_NN_ and every figure quoted above belongs to *d*_NN_ = 2.4 voxels. At the working deck *d*_NN_ = 1.47 voxels and recall is exactly 1.0000 on all four systems, so there the floor lies below the smallest component present in any of them. The floor was not re-measured at the working deck.

Figure 4(b) shows the floor operating on the two-cluster E5 system, at a 3.5 Å grid spacing. Two clusters of 22 voxels are separated by a 20 Å gap. At ALPHA = 3.5, hence *d*_NN_ = 8.4 Å, the lattice places no seed on either cluster and BLS returns an empty set. At ALPHA ≤ 3.2 seeds land on both and both are recovered, which includes the working value of 2.45. The single dip at ALPHA = 3.0, where four seeds are placed and one lands on a cluster, shows that the floor depends on where the lattice phase falls as well as on its period, which is why the 192-cell matrix sweeps eight origin offsets.

### 4.5 Detection Limits

Percolation and gap resolution, both bound what can be recovered, and both are properties of the discretized problem. Both were measured at a 3.5 Å grid spacing.

#### Percolation

In E4 the cluster count rises in a fixed volume (Figure 5a). The number of physically separate objects tracks the nominal count exactly through *N* = 150, falls to 178 of 200 and 212 of 250, then to 5 at *N* = 300 and 1 at *N* = 750. The onset bracket is therefore *N* = 250 → 300, located from the data as the adjacent pair with the largest fractional collapse, a factor of 42.4 against a smaller change for every other pair. The largest connected component goes from 116 voxels at *N* = 250 to 10 606 at *N* = 300, a factor of 91: a spanning cluster has formed [14] and the nominally distinct clusters have physically merged. Partial merging begins at *N* = 200, where 11.0% of the nominal clusters are already joined. Past the onset there are no longer *N* separable objects to find, so recall against the expected count stops being a meaningful metric. Every exact method returns identical numbers here, which confirms the merging. E3 is the control: when the volume grows with the cluster count so that density stays fixed, detection remains complete out to 5000 clusters. Recall in both experiments divides by the DFS-measured present count, which differs from the packing request wherever overlaps merge clusters: 500 requested yields 499 present, 1000 yields 995, 2000 yields 1986 and 5000 yields 4862.

**Figure 5:**
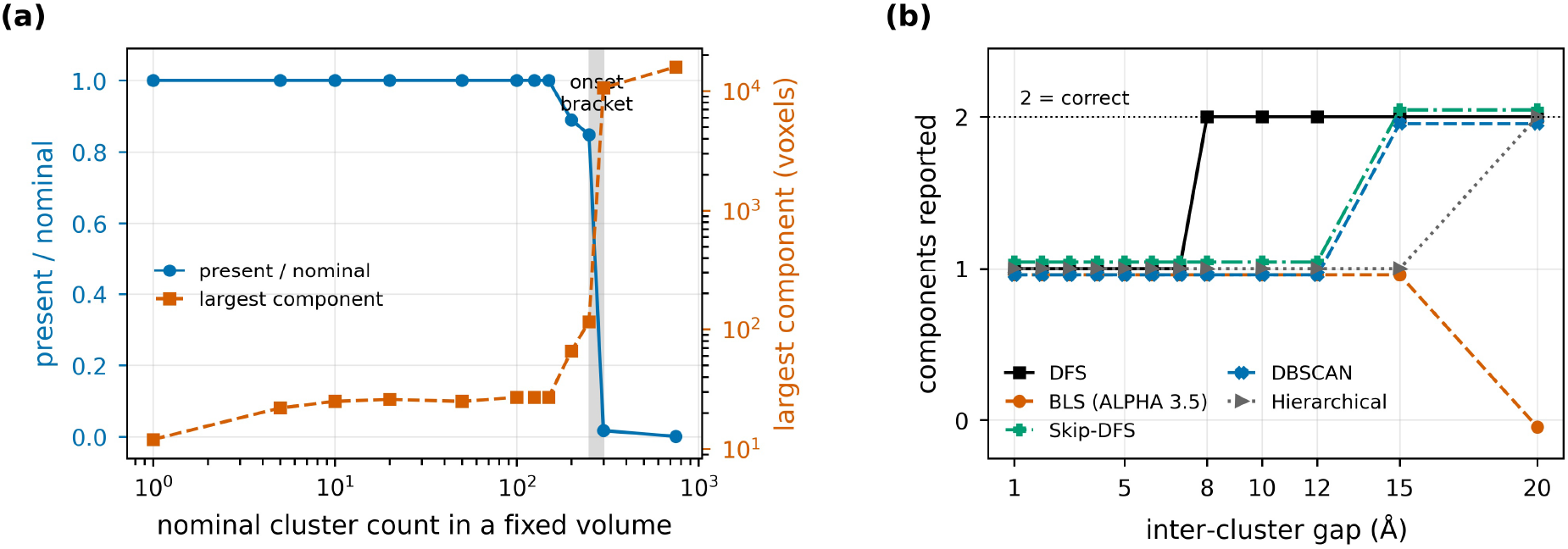
Percolation and gap resolution. Two limits on what any connectivity-based method can recover, both at a 3.5 Å grid spacing. (a) Percolation onset (E4 experiment), cluster count increases in a fixed volume. The number of separate objects present divides by the nominal count on the left axis; the largest connected component is on the right. The shaded band is the onset bracket located from the data. The count falls because clusters join, as the 91-fold jump in the largest component shows. (b) Gap resolution (E5). Exhaustive methods step from one component to two at 8 Å; Skip-DFS and DBSCAN at 15 Å; agglomerative clustering at 20 Å. The BLS curve is at ALPHA = 3.5, at which the 22-voxel clusters lie below its size floor. Coincident curves are drawn with a small vertical offset so that both remain visible.

#### Gap resolution

In E5 two clusters are separated by a controlled gap (Figure 5b). Every exhaustive method resolves them at 8 Å and fails at 7 Å, a step. The threshold is set by the grid resolution and the cutoff and is shared exactly by all of them, so it belongs to the discretization. Skip-DFS at unit stride and DBSCAN require 15 Å, agglomerative clustering 20 Å. BLS was run on E5 at ALPHA = 3.5 and reports one component from 1 Å to 15 Å and none at 20 Å, because the 22-voxel clusters lie below the floor that value of ALPHA sets. The sweep of Figure 4(b) recovers both clusters at the same gap at the working ALPHA.

### 4.6 Strided Traversal

A variation of the DFS algorithm, was implemented. Skip-DFS with a stride *s >* 1 was included to quantify what edge verification is worth. Every stride is slower than DFS, at 1.042 to 1.099 times its cost, so there is no speed-up against which to trade accuracy. Accuracy and resolution degrade monotonically with *s*: the cluster count falls from 1.000 to 0.985 of the DFS count while the maximum cluster size rises from 1.000 to 1.362, and the gap needed to resolve two clusters grows from 8 Å at *s* = 1 to 20 Å at *s* = 4, with *s* = 5 never resolving within the tested range. The two ratios moving in opposite directions identifies the mechanism: the stride merges components that the exact tier separates. Only *s* = 1 preserves accuracy and resolution, and it is still 4.2% slower than plain DFS. There is no stride at which strided traversal is worth its cost on these systems, and BLS uses unit stride throughout. The full stride table is given in the Supplementary Information.

### 4.7 Probe-Budget-Normalized Lattice Comparison

Everything reported above holds *d*_NN_ fixed when varying the lattice, which gives the centerings unequal probe budgets. The discriminating experiment fixes the probe count and sweeps the cluster diameter downward through *d*_NN_, recording for each of P, I and F the diameter at which recall first departs from 1.0. If the covering-radius account is correct, those thresholds should stand in the ratio 1.859: 1.000: 1.431 for P, I and F, the cubes of Equation (4) relative to the body-centred lattice. The criterion was fixed before the experiment ran: confirmation would make the lattice a design variable with a computable optimum and would imply that the face-centred default is the wrong one; a null result would mean the crystallographic framing contributes nothing beyond *d*_NN_-scaled spacing.

The outcome is partial (Table 5, Figure 6). Three components separate.

**Table 5:** Probe-budget-normalized lattice comparison (E6). Predicted ratios are the cost-normalized covering radii of Equation (4) cubed, relative to the body-centred lattice. Measured ranges span six recall thresholds. The decision rule was fixed before the experiment ran: confirmed if the ratios match within the budget-equalisation tolerance (3.42%), null if no ordering survives, partial otherwise. Cross-arm agreement is 3.7% maximum deviation over nine pairings. Overall verdict: partial.

| Component | Predicted | Primary arm | Control arm | Outcome |
| --- | --- | --- | --- | --- |
| P/I ordering | 1.859 | 1.066–1.294 | 1.015–1.655 | survives |
| P exceeds I in: primary arm 6/6; control arm 6/6; hexagonal cells 6/6; triclinic cells 11/12. |  |  |  |  |
| P/I magnitude | 1.859 | 1.066–1.294 | 1.015–1.655 | not recovered |
| 0/12 pooled values within tolerance; shortfall 30–43% (primary), 11–45% (control). |  |  |  |  |
| F/I | 1.431 | 0.885–1.069 | 1.000–1.177 | unsupported |
| 0/12 within tolerance of the prediction, but 10/12 within 10% of unity. |  |  |  |  |

**Figure 6:**
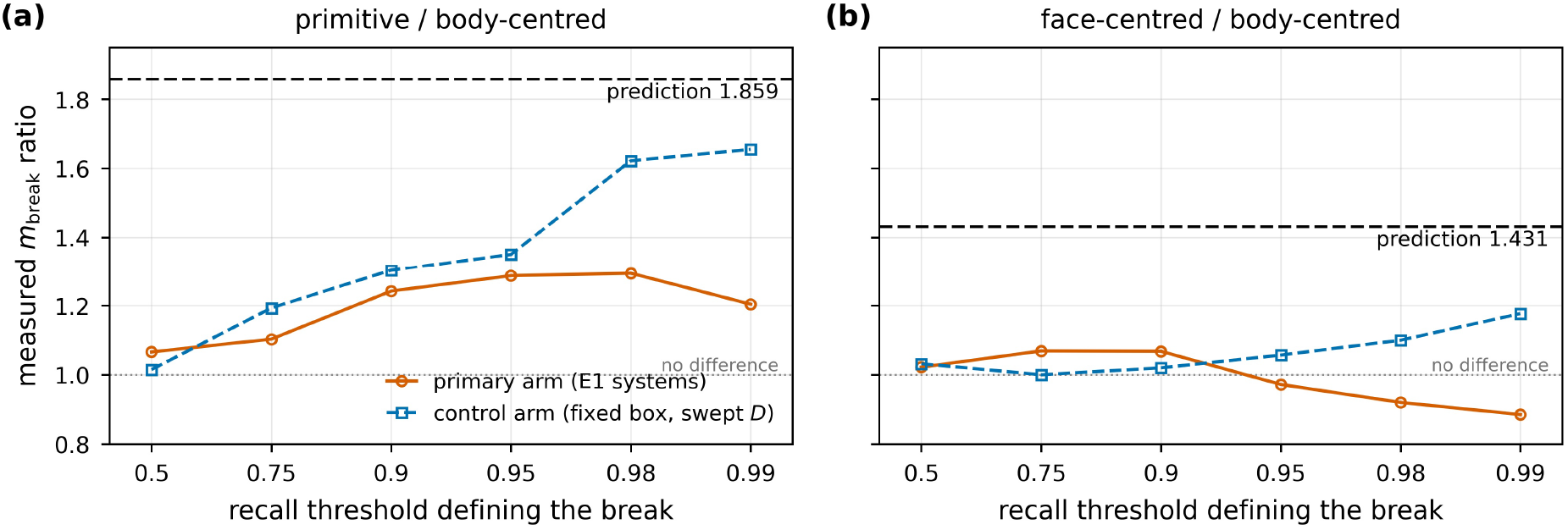
Probe-budget-normalized comparison of the three cubic centerings (E6), for (a) primitive against body-centred and (b) face-centred against body-centred. Each panel shows both arms against the prediction of Equation (4) cubed. The primitive-to-body-centred ordering survives in direction in both arms and reaches at most 70% of the predicted magnitude, rising as the recall threshold tightens. Face-centred and body-centred are indistinguishable at equal probe count.

#### The ordering survives

P exceeds I at all six pooled recall thresholds in both arms, in 6 of 6 hexagonal cells and in 11 of 12 triclinic cells. The direction of the Bambah and Conway–Sloane result [12, 13], that the body-centred lattice is the thinnest lattice covering of three-space, holds in the measurement.

#### The magnitude falls short

Against a prediction of 1.859 the primary arm measures 1.066–1.294 and the control arm 1.015–1.655, shortfalls of 30–43% and 11–45% respectively. None of the twelve pooled values falls within the budget-equalisation tolerance of the prediction.

#### F and I are indistinguishable at equal probe budget

Against a prediction of 1.431 the primary arm measures 0.885–1.069, mean 0.989, and the control arm 1.000–1.177, mean 1.064. Ten of the twelve pooled values fall within 10% of unity. The face-centred default therefore carries no measurable probe-cost penalty, and the earlier claim that a body-centred default would achive a 1.43-fold reduction in probes is withdrawn. The centering check of Figure 2(b) reaches the same conclusion at the working deck by a different route.

The two arms agree with each other and both disagree with the prediction. Cross-arm agreement is 3.7% at worst and 1.5% on average over nine threshold-by-centering pairings, with all nine within 5%, while the arms look at different clusters: the control arm sweeps effective diameters of 4.29 to 10.97 voxels and the primary arm’s marginal cluster sits at 3.3 to 3.9. The agreement is a genuine cross-check.

The compression has a straightforward explanation. The covering radius is a worst-case quantity, the radius of the largest empty hole, whereas recall over a population of clusters is an average. At equal density all three centerings cover a typical cluster alike and differ only on the rare worst-placed one. The P/I ratio rising from 1.07 to 1.29 as the threshold tightens from 0.50 to 0.98 is the signature of an effect that lives in the tail of the distribution. Timing is flat across centerings to within 1% on every system.

The quality of budget equalisation is set by how many lattice periods span the box: 14 periods gave a 10.8% spread in achieved probe count, while 105 periods gave 1.6%. The primary arm reaches 0.92% median spread because it runs at 294^3^.

## 5 Discussion

Conventional CCL finds its seeds by rastering the grid. BLS instead evaluates a sparse probe set, spaced by a derived covering radius, and expands from each occupied probe by exact traversal. Sparse seeding and depth-first expansion are not new individually; the pairing is: a probe spacing that is derived rather than chosen, sparse enough to remove the raster scan, and specific enough to state in advance what it can miss.

BLS returns exact connected components above a stated size floor: over 192 cells nothing was truncated, split or merged, so its output is always a subset of the true component set. The floor is a query parameter, as the deck search of Section 4.1 shows. Requiring recall 1.000 on every system, the fastest admissible *d*_NN_ sits below the smallest component present, and BLS then returns the complete component set faster than any exhaustive method compared here. In nucleation analysis the floor is itself the quantity of interest, since sub-critical nuclei are discarded anyway; coarsening *d*_NN_ further trades speed against what is returned.

Bravais lattices provides a limited improvement: E6 confirms the predicted ordering of centerings at equal probe budget, but not the predicted magnitudes, and *F* and *I* are indistinguishable at equal cost. It still explains why *d*_NN_-scaled spacing works and what sets the floor. Cubic-*F* remains the default and suffices for all four morphologies tested, but lattice and centering stay the most consequential choice a user makes: they span 0.21 and 0.12 in recall against at most 0.009 for every other parameter, and cubic-*P* already fails the recall constraint at the working deck.

On cost, BLS is a DFS-class method. The tier it shares with DFS comes from traversal proportional to occupancy, which is a property of depth-first expansion. The lattice contributes the seed scan, and at the working deck that scan examines 44.5% of the sites a raster scan would, for a 4–8% saving in total runtime. The saving is smaller than the reduction in site count, and the margin is comparable to the replicate spread on two of the four systems.

Limitations of BLS and the current work: Memory cost for BLS is *O* (*M* ^3^) for the grid itself. The floor guarantee is one-sided and was measured only on randomly packed clusters; oriented or commensurate structures could interact with the lattice phase and remain untested. No periodic boundary handling exists, so all results are for raw voxel adjacency. Variations of cloud shape are provided, but not confinement geometry, since its cell stays orthorhombic throughout. At fine grid spacings the disordered system over-segments, hence recall for those systems is reported as a minimum across systems. The size floor and the two detection limits were measured only at 3.5 Å and not re-checked at the working deck.

The construction assumes only a binary occupancy field on a regular grid and a target size known in advance, so it transfers wherever those hold and the occupied fraction is small. Digital rock physics, volumetric medical segmentation, industrial computed tomography and halo finding in cosmological simulations all satisfy the first two conditions. Whether the probe saving is worth having depends on the third, and on the ratio of grid volume to target size. We have not tested any of these domains and make no claim beyond the structural one.

The two limits of Section 4.5 belong to no method in particular. Past the percolation threshold the useful question becomes how large the connected network is, and exact CCL gives a clean criterion for locating that crossover.

## 6 Conclusions

- BLS replaces the raster seed scan of conventional CCL with a sparse lattice probe scan and returns exact connected components, subject to a size floor that follows from the covering radius. The pairing of sparse seeding with an advance statement of what can be missed is what distinguishes it from uniform-grid seed-and-grow methods.
- At the working deck, 4.0 Å grid spacing with ALPHA = 2.45 and a cubic-*F* probe lattice, BLS reaches recall 1.0000 on all four ice morphologies and is the fastest method in the comparison, at 0.917–0.958 times DFS. CC3D and RLE-CCL cost 1.06–1.16 times DFS and GCBD 2.59–3.43 times. A single deck serves all four systems.
- On every component it reaches, BLS agrees with an exhaustive labeler exactly: nothing truncated, split or merged in 192 of 192 cells spanning four systems, six lattices and eight lattice-origin offsets. Its output is a subset of the true component set.
- Lattice and centering are the parameters that matter, spanning 0.21 and 0.12 in recall against at most 0.009 for every other configurable parameter. Cubic-*P* fails the recall constraint at the working deck; cubic-*I* and cubic-*F* both meet it.
- The size floor measured for the tested systems at *d*_NN_ = 2.4 voxels is 60 voxels pooled over all six lattices tested and 29 for the cubic-*F* default alone.The floor scales with *d*_NN_, and setting *d*_NN_ below the smallest component of interest holds the guarantee empirically.
- Grid spacing couples runtime to recall. Every timing in this paper states its spacing and transfers to another only on re-measurement. The relative margin over DFS narrows from 0.81–0.86 at 2.5 Å to 0.91–0.94 at 4.0 Å while absolute runtime falls throughout.
- The speed advantage comes from removing the raster seed scan, which at the working deck examines 44.5% of the sites a full scan would. The resulting saving in total runtime is 4–8%, comparable to the replicate spread on two of the four systems.
- Skipping cells during traversal is not a useful approximation on voxel grids at these occupancies. Every stride from 2 to 5 is both slower than DFS and less accurate, and the stride merges components.
- At equalized probe budget the predicted ordering of the centerings holds, but its magnitude does not, and the face-centred and body-centred lattices are indistinguishable. The face-centred default stands and the 1.43-fold probe-cost does not hold.
- Percolation and gap resolution bound any connectivity method, not just this one.

## Author contributions

Conceptualization, F.C.; writing, original draft preparation, F.C.; data curation, F.C.; review and editing, F.C. The author has read and agreed to the published version of the manuscript.

## Funding

This research was supported by grant 0311/SBAD/0781 from Poznan University of Technology. Selected computational results reported herein were independently verified at the Poznan Supercomputing and Networking Centre.

## Data availability

https://github.com/franciscocarrascoza/BLS

## Acknowledgments

The infrastructure of the European Centre for Bioinformatics and Genomics affiliated with the Poznan University of Technology was used in this work.

Computations to corroborate some of the information presented were performed at the Poznan Supercomputing and Networking Centre. Computational code was developed with the assistance of a generative artificial-intelligence tool (Claude Opus 5, Anthropic); the extracted values were checked against the source by the author, who takes full responsibility for their accuracy.

## Conflicts of interest

The author declares no conflicts of interest.

## Notes

https://github.com/franciscocarrascoza/BLS

### Competing Interest Statement

The authors have declared no competing interest.

